# To fly, or not to fly, that is the question: A deep learning model for peptide detectability prediction in mass spectrometry

**DOI:** 10.1101/2024.10.28.620610

**Authors:** Naim Abdul-Khalek, Mario Picciani, Reinhard Wimmer, Michael Toft Overgaard, Mathias Wilhelm, Simon Gregersen Echers

## Abstract

Identifying detectable peptides, known as flyers, is key in mass spectrometry-based proteomics. Peptide detectability is strongly related with the peptide sequence and its resulting physicochemical properties. Moreover, the high variability in MS data, particularly in peptide detectability and intensity across multiple analyses and samples, makes the development of a generic model for detectability prediction unfeasible. This underlines the need for tools that can be refined for specific experimental conditions. To address this need, we present Pfly, a deep learning model developed to predicts peptide detectability based solely on peptide sequence. Pfly distinguishes itself as a versatile and reliable state-of-the-art tool, offering high performance, accessibility, and easy customizability for end-users. This adaptability allows researchers to tailor the model to their specific experimental conditions, facilitating the creation of lab-specific models. This, in turn, can lead to more accurate results and expand the model’s applicability across various research fields. The model’s architecture is an encoder-decoder with an attention mechanism. This tool classifies peptides as either flyers or non-flyers, providing both binary probabilities and detailed categorical probabilities for four distinct classes defined in this study: non-flyer, weak flyer, intermediate flyer, and strong flyer. The model was initially trained on a synthetic peptide library and subsequently fine-tuned with a biological dataset to mitigate bias towards synthesizability, improving the predictive capacity and outperforming state-of-the-art predictors in a benchmark comparison. The study further investigates the influence of protein abundance and the search engine, illustrating the negative impact on peptide identification due to misclassification. Pfly has been integrated in the DLOmix framework and it is accessible on GitHub at https://github.com/wilhelm-lab/dlomix.

## 1. Introduction

Mass Spectrometry (MS) stands as a cornerstone in proteomics due to its sensitivity, selectivity, and high throughput capabilities, pivotal across academic and industrial sectors such as pharmaceuticals, medicine, and food. Although MS technology has improved considerably over the years, drawbacks and limitations remain, leading to variability in the results obtained from LC-ESI-MS analysis^1–6^. This variability is often reflected in the inconsistent detectability and intensity of peptides, where more abundant peptides can suppress the detection of lower-abundance ones, and differences in ionization efficiency further contribute to fluctuating intensity responses. Within the sources of variability, one significant factor is sample complexity. Biological samples often contain a large number of molecules (e.g. proteins) with a wide dynamic range of concentrations, which can negatively impact identification and subsequent quantification of individual proteins and peptides^7^. Moreover, variability in MS analysis is influenced by instrumental design^6–11^ and downstream processes, such as the databases used for peptide and protein identity inference and the selected search engine, which can further exacerbate differences in the results^8,12^.

A significant challenge in MS-based proteomics is peptide detectability, as not all peptides present in the sample are detected, and even those present in equimolar amounts may exhibit different signal responses^13^. Ionization efficiency, influenced by peptide sequence and its physicochemical properties such as hydrophobicity, significantly contributes to this variability^14–16^. During reverse phase high-performance liquid chromatography (RP-HPLC), more hydrophobic peptides are retained more efficiently on the column and elute at a higher concentration of the organic solvent, usually acetonitrile, in the mobile phase. This organic solvent evaporates more readily than water, facilitating a more efficient gas-phase transition of the peptides^8,17^. Additionally, hydrophobic peptides also tend to partition into the organic phase and remain closer to the droplet surface during electrospray ionization (ESI), which further facilitates their transition to the gas phase^14,16,18^. On the other hand, the presence of positively charged amino acids (AAs), namely arginine (Arg), lysine (Lys), and histidine (His) within the peptide sequences, increases the likelihood of obtaining positively charged ions^19,20^, critical for MS detection and robust signal response in positive mode analysis^8,17,21^. Moreover, aspects such as protein/peptide abundance^22–24^ as well as co-elution^15^, competition for ionization^22^, ion suppression^15,25^, and co-fragmentation^26^ have also been emphasized for their effects on peptide detectability.

Despite this complexity, the pursuit of an accurate peptide detectability model remains crucial for advancing proteomic research and its applications. Detectability prediction could impact many aspects of MS-based proteomics through integration into quality assessments of search engine identifications^27^. This may be a valuable tool for identification of proteins that are typically missed due to their proteotypic peptides (PTPs) all having low detectability, thus presenting an inherent barrier to identification via MS-proteomic analysis. Moreover, in large-scale proteomic studies, accurate prediction models can optimize experimental workflows by prioritizing peptides likely to be detected, thereby streamlining data acquisition and analysis processes^22,28,29^. Specifically, these models are invaluable for designing targeted MS analyses, relying on accurate prediction of peptide detectability to focus on the most promising candidates^30,31^. Additionally, peptide detectability has previously been used for spectral library filtering in data-independent acquisition (DIA) by limiting the MS2 fragmentation spectrum search space for search engines, leading to higher efficiency and accuracy of peptide identification^32^. This capability is crucial across diverse fields, including personalized medicine and clinical research as it enables more precise biomarker discovery, which in turn can facilitate the development of diagnostic tools and therapeutic targets^33–35^. In the realm of food science, robust peptide detectability models can be of great aid for ensuring food safety and quality assurance^36–39^. They can assist in the identification of allergens, contaminants, and authenticate food products, thus strengthening consumer trust and regulatory compliance. By overcoming the inherent challenges of variability in MS-based proteomics, accurate peptide detectability models will pave the way for more reliable and reproducible research outcomes across diverse applications in many fields such as biology, medicine, and food science.

Taken together, the specific characteristics and properties of analytes, along with the complexity of sample composition and analytical conditions, pose a formidable challenge in developing a universal model for peptide detectability in MS. This challenge is compounded by the variability introduced by the diverse MS technologies used across laboratories, encompassing differences in instrumental design, data acquisition methods, sample preparation protocols, and quality control procedures^40–43^. All these factors collectively contribute to the significant variability in MS-based proteomic analyses, hindering the establishment of a one-size-fits-all approach to peptide detectability prediction.

Using AI for predicting peptide detectability is not a new concept, as several published studies introduce computational tools for this purpose. Notable models include PeptideSieve^44^, CONSeQuence^45^, PPA^24^, d::pPop^46^, AP3^47^, DeepMSPeptide^48^, PepFormer^23^, CapsNet-CBAM^49^, DeepDetect^32^, and DbyDeep^50^. These tools are accessible either through web platforms, downloadable software packages, or provided as source code. However, many of these tools are not trivial to use, as either the web-tools are currently not working or modifications to the source code are necessary, thereby significantly reducing accessibility and applicability. Several tools are designed to predict detectability based on computed peptide-level properties, thereby overlooking sequence-specific effects and potentially affecting the models’ performance. Moreover, due to the complexity of the peptide detectability prediction task, it is highly beneficial to facilitate easy model fine-tuning with lab-specific data to reduce inherent biases. This refinement is necessary to enhance performance for specific applications and to generate lab-specific models that would outperform more generic models. Unfortunately, all the mentioned tools lack a simple and user-friendly pipeline for customization, thereby restricting users with limited or no programming skills from fully leveraging a model’s predictive capabilities.

Thus, there is a clear need for a state-of-the-art peptide detectability model that is not only freely and easily accessible to the proteomics community but also provides a reliable platform with an intuitive and customizable pipeline. This platform should allow users to easily use, train, and fine-tune the model with their specific data. Such a solution would bridge the current gap, enabling more researchers to benefit from high-performance peptide detectability models without the barrier of technical complexity. To address this need, we present “Pfly”; a customizable deep learning model for peptide detectability prediction.

Pfly is an encoder-decoder with an attention mechanism, designed as a multi-class model but modified to generate a binary output requiring only peptide sequences as input. Pfly was initially trained on an intended equimolar, synthetic peptide library using sequence and intensity information from LC-ESI-MS analysis. It was subsequently fine-tuned using a large dataset of enzymatically digested proteomes from different human cell lines. The predictive capabilities of Pfly were validated through benchmarking against state-of-the-art detectability predictors, including PepFormer^23^, DeepMSPeptide^48^, and DeepDetect^32^. Pfly was incorporated into DLOmix^51^, a Python framework for deep learning (DL) in proteomics. Within DLOmix, the pipelines are provided for direct use, training, and fine-tuning. The source code is freely available, empowering users with varying levels of programming skills to utilize this tool effectively and more experienced programmers to expand upon its capabilities. Pfly can support diverse applications, including precursor peptide selection for targeted MS analysis, experimental workflow optimization, improved peptide identification accuracy, and facilitate reliable research across various fields.

## 2. Materials and Methods

### 2.1. Development and initial evaluation dataset

An initial model, the base model, was trained and tested using the ProteomeTools data extracted from the PRIDE repository with the identifiers <u>PXD004732</u>, <u>PXD010595</u>, and <u>PXD021013</u>, and the supplementary material from the corresponding publications^52–54^. The datasets include over 1 million unique synthetic peptide sequences selected from the human proteome.

For flyer peptides (i.e., experimentally identified), data filtering was carried out (Fig. S1) similarly to previous studies^17,55^. The intention is to improve the quality of the data to build the most optimal model for peptide detectability. Initially, peptides with a PEP score equal or higher than 0.01 were removed, as well as reverse order sequences, potential contaminants, and peptides with no intensity. Then, peptides that were identified by using a specific enzyme and with trypsin as proteases were selected, removing all duplicated sequences, and keeping the median values of the MS1 intensity output. Finally, the coefficient of variation (CV) was computed for recurring sequences, removing those with a CV higher than 0.54, effectively preserving essential content while mitigating high variability. After this filtering process, the flyer dataset consisted of 251,070 unique peptides. This dataset was ordered based on MS1 intensity values and separated into three equal quantiles. Peptides in the low-intensity tertile were classified as weak flyers, those in the center tertile as intermediate flyers, and those in the high-intensity tertile as strong flyers (Fig. 1A).

**Figure 1.**
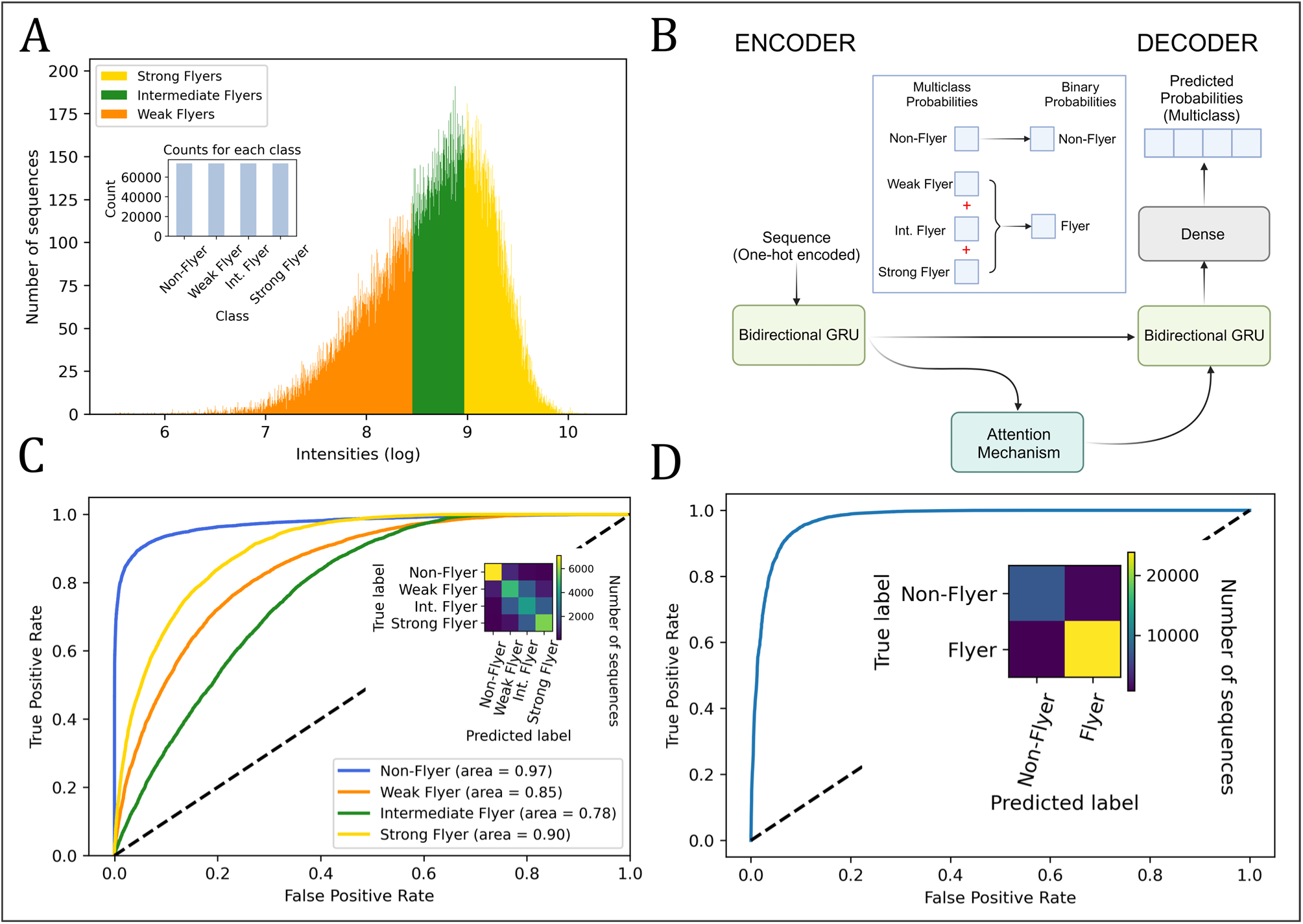
Building and testing an encoder-decoder with an attention mechanism for predicting peptide detectability. (A) Data splitting and labeling based on peptide MS1 response. The ProteomeTools data was divided into tertiles (three quantiles of equal size) along the log intensity dimension to separate flyers into three classes representing weak flyers (orange), intermediate flyers (green), and strong flyers (yellow), according to their MS1 intensity. Non-flyers were the peptides that were synthesized but not observed in the experiments. The dataset was balanced through downsampling to ensure equal representation of all classes (see inset) (B) Schematic representation of the model architecture. The model architecture is an encoder-decoder enhanced by an attention mechanism. The encoder features a single bidirectional GRU layer responsible for encoding input sequences into a new representation. The attention mechanism processes the encoder’s output, which is then fed into the decoder. The decoder consists of a bidirectional GRU layer and a dense layer with a softmax function. This dense layer comprises 4 units, each representing the probability of one of the four classes: non-flyer, weak flyer, intermediate flyer, and strong flyer. The predicted probabilities for all flyer classes (weak, intermediate, and strong) are aggregated to compute the probability of a peptide being a flyer, while the probability for non-flyers remains unchanged. A more detailed schematic of the model architecture can be found in the supplementary information (Fig. S4). (C) Performance of the base model for categorical predictions on the ProteomeTools test dataset. ROC curves show the model’s ability to correctly predict the labels for non-flyers (blue), weak flyers (orange), intermediate flyers (green) and strong flyers (yellow), along with the area under the curve for each class. Inset: Confusion matrix of the multiclass results, illustrating the base model’s performance across all classes and its ability to discriminate between them, with counts ranging from low (dark blue) to high (yellow). (D) Performance of the base model for binary predictions on the ProteomeTools test dataset. The ROC curve shows the model’s ability to discriminate between flyers and non-flyers (area = 0.97). Inset: Confusion matrix for binary prediction, with counts ranging from low (dark blue) to high (yellow).

The non-flyers (i.e., not experimentally identified), on the other hand, were obtained from the full list of synthetized peptides in the original studies and identified by discarding the peptides that were observed during the MS analysis.^53,54^ Duplicated sequences were removed as well as sequences containing other than the 20 standard AAs. In total, the non-flyer dataset consisted of 82,664 unique peptides.

Afterwards, the flyer and non-flyer datasets were merged, duplicated peptides between flyers and non-flyers were identified and removed, and the merged dataset was downsampled based on the minority class (i.e. non-flyer peptides). The final dataset consisted of 328,832 unique tryptic peptide sequences. This dataset was randomly split into subsets for training and testing (90:10). From the training set, 20% was randomly sampled to be used as the validation set.

### 2.2. Refinement and evaluation dataset

The dataset deposited in the ProteomeXchange Consortium via the MAssIVE partner repository with the identifier <u>PXD024364</u>, as described by Sinitcyn *et al.* ^56^, was used to fine-tune the base model and reduce bias towards synthesizability, resulting in the final model termed “Pfly”. This refinement was performed using one subset of this dataset, while the other subset was reserved for evaluating Pfly, the base model, and other tools. This dataset contains bottom-up proteomics data from six different human cell lines (GM12878, HeLa S3, HepG2, hES1, HUVEC, and K562), deep fractioning (24–80 fractions) and three different fragmentation methods (HCD, CAD and ETD). All cell lines were digested with six different proteases (LysC, LysN, AspN, chymotrypsin, GluC and trypsin). From all proteins identified in this dataset, only the uniquely identified canonical proteins sequences (UniProtKB/Swiss-Prot) were considered. The mean and standard deviation of the proteins’ iBAQ intensity responses were computed from the cell lines in which they were identified using tryptic digestion.

After identifying a total of 1,133,869 peptides, a filtering step was implemented to refine the dataset (Fig. S2). Specifically, 4,526 peptides were eliminated due to being reverse sequences, 9,833 peptides were identified as potential contaminants, and an additional 96,283 peptides were excluded because they either did not originate from any canonical proteins or could belong to more than one (i.e. razor peptides). After this initial filtering, 1,023,227 peptides remained, originating from a total of 12,333 unique canonical proteins. Additionally, a feature (defined as flyability) was generated, using the number of cell lines digested with trypsin from where a particular peptide was identified. For example, if a peptide was quantitatively identified in 5 of the 6 cell lines, it would be assigned a flyability value of 5/6 (0.833). Next, an *in silico* tryptic digest (using pyteomics^57^) was performed on the 12,333 proteins, with a minimum peptide length of 7 AAs and no missed cleavages. From the resulting peptides, duplicates were filtered out, leading to a total of 374,160 unique tryptic peptides. If any of those peptides were found in the list of experimentally identified peptides and had a flyability score greater than zero, they were categorized as flyers; otherwise, they were considered non-flyers. Flyers were further classified based on their flyability scores: Peptides with scores between 0 and 0.5 were considered weak flyers, those with scores between 0.5 (inclusive) and 0.8 were classified as intermediate flyers, and those with scores greater than or equal to 0.8 were categorized as strong flyers. Labels were also assigned based on the intensity responses within individual cell lines digested with trypsin. If a peptide showed an intensity response in a particular cell line, it retained its previously assigned labels; if not, it was categorized as a non-flyer. The 374,160 peptides that resulted from the *in silico* tryptic digest underwent further filtering. Specifically, 14,257 peptides were eliminated due to their length exceeding 40 AAs, and an additional 22 peptides were excluded because they contained other than the 20 standard AAs.

The dataset was reduced to 359,881 peptides assigned to 12,325 canonical proteins. For the test dataset, 2000 proteins were randomly selected. These proteins had iBAQ values greater than zero for the randomly selected GM12878 cell line, which was digested with trypsin. The test proteins represent 60,185 peptides from which 40,846 were flyers and 19,339 were non-flyers. The training/validation dataset consisted of the remaining 10,325 proteins. This dataset included 299,696 unique peptides, of which 183,489 were flyers and 116,207 were non-flyers. To balance the classes, the dataset was randomly downsampled based on the minority class, reducing each class to 46,634 peptides. Ultimately, the final training/validation dataset consisted of 186,536 peptides assigned to 10,285 canonical proteins, split with a 90:10 ratio between training and validation sets, respectively.

### 2.3. Final evaluation dataset

The dataset described by Wang *et al.*^58^, deposited in the ProteomeXchange Consortium via the PRIDE partner repository with the identifier <u>PXD010154</u>, was used for a final evaluation and performance benchmarking. This particular dataset was not used for training or fine-tuning; thus, it provided a more unbiased evaluation of the fine-tuned model (Pfly), the base model, and the other tools. This dataset represents 30 healthy tissues from the Human Protein Atlas Project. The proteins identified in this study were considered if they could be successfully mapped from Ensembl to a unique canonical protein (using UniProt/Swiss-Prot as reference). Next, the iBAQ mean and standard deviation were computed using the data from all tissues where the proteins were identified.

A total of 277,698 peptides were identified. An initial filtering process was conducted (Fig. S3), removing 1,885 peptides identified in the reverse order, 2,145 potential contaminants, and 114,840 peptides that did not originate from mapped unique canonical proteins. After the initial filtering, the total number of peptides dropped to 158,828, originating from 7,389 proteins. Similar to the dataset used for fine-tuning, peptide flyability was defined based on the number of tissues in which each peptide was identified with an intensity greater than zero. For example, if a peptide had an intensity response in 29 tissues, the assigned flyability score would be 29/30 (0.967). Afterwards, an *in silico* tryptic digest (using pyteomics^57^) was performed on the 7,389 Ensembl proteins, allowing a minimum peptide length of 7 AAs and no missed cleavages. After removal of duplicate peptides, the list comprised 255,358 tryptic peptides. If a peptide was found in the list of experimentally identified peptides and had a flyability score greater than zero, it was classified as a flyer; otherwise, it was considered a non-flyer. Flyers were further categorized based on their flyability scores: Peptides with scores between 0 and 0.333 (inclusive) were considered weak flyers, those between 0.333 and 0.7 were defined as intermediate flyers, and those at or above 0.7 were classified as strong flyers. Additionally, labels were assigned according to intensity responses in specific tissues. If a peptide had an intensity response in a particular tissue, it retained its previously assigned labels; if it did not, it was labeled as a non-flyer. The 255,358 tryptic peptides were further filtered, removing 9,851 peptides exceeding 40 AAs in length and 7 peptides containing symbols that do not correspond to any of the 20 standard AAs. The dataset was reduced to 245,500 peptides assigned to 7,371 canonical proteins, from which only the proteins with iBAQ intensities higher than zero for the randomly selected tonsil tissue (reference id P010747), were considered. Thus, the total number of proteins used for testing was 5,565. The test proteins amounted to 190,955 peptides whereof 88,197 are classified as flyers and 102,758 as non-flyers.

### 2.4. Model Architecture

The models’ architecture is an encoder-decoder with an attention mechanism (Fig. 1B, S4), which is based on bidirectional recurrent neural networks (BRNN)^59^ with gated recurrent units (GRU)^60^. The encoder consists of one bidirectional GRU layer. The decoder has the same configuration as the encoder but with the addition of a dense layer with a softmax function. The dense layer has 4 units, each representing the probability of one of the four classes: non-, weak, intermediate, and strong flyer. The prediction is determined by selecting the class with the highest probability; if any of the flyer classes has the highest probability, the peptides are labeled as flyer; otherwise, they are classified as non-flyer. The binary probabilities (flyer/non-flyer) are computed by summing the predicted probabilities of the flyer classes (weak, intermediate, and strong) while the predicted probability for the non-flyer class remains untouched. The model input has a dimension of 40 x 21, representing one-hot encoded peptide sequences with a maximum length of 40 AAs, where padding is applied for shorter peptides. Thus, there are 21 tokens: the 20 natural AAs in addition to the padding character. The number of units for the BRNN with GRU is 64 (for the encoder and decoder) and the batch size is 128.

### 2.5. Training and testing

The models and all the results were obtained using Python (v.3.8.8) with TensorFlow^61^ (v. 2.5.0) and the following libraries: Scikit-learn^62^ (v.1.1.2), Statsmodels^63^ (v.0.13.2), Pandas^64^ (v.1.4.4), Matplotlib^65^ (v.3.5.2), Seaborn^66^ (v.0.11.2), SciPy^67^ (v.1.9.1), and NumPy^68^ (v.1.23.1), pyteomics^57^ (v.4.6.3).

Categorical cross entropy was used to calculate the loss and categorical accuracy was used to measure the model performance on the validation dataset during training. The final results were evaluated through various methods and associated metrics, including binary accuracy, precision, recall, F1 score, Matthews’s correlation coefficient (MCC), area under the curve (AUC), receiver operating characteristic curve (ROC curve), and confusion matrix. Adam^69^ was the optimizer chosen. The models were trained on NVIDIA Quadro T2000 GPU. The model was developed using the ProteomeTools data (see Methods). During training, overfitting was controlled using the validation data, while the test data was used to do an initial evaluation of the base model and its generalization capabilities. The model was optimized using different hyperparameter settings, e.g., batch size, number of epochs, number of units per layer, numbers of layers.

The Sinitcyn *et al.* dataset^56^ (see Methods) was used to fine-tune the base model. A subset from this dataset was also used to test the base model, the fine-tuned (Pfly) model, PepFormer^23^, DeepMSPeptide^48^, and DeepDetect^32^, allowing a performance comparison between them. The Wang *et al.* dataset^58^ (see Methods) was exclusively used for a final evaluation of the base and final (Pfly) model as well as the other tools in a benchmark comparison. Lastly, the Sinitcyn *et al.* test dataset was rescored using Oktoberfest^70^ (v.0.6.2), and the performance of Pfly was compared between the rescored and original test dataset.

Source code and scripts for Pfly have been integrated in the DLOmix framework and it is accessible on GitHub at https://github.com/wilhelm-lab/dlomix.

## 3. Results and Discussion

For the development and initial evaluation dataset(see Methods), an initial data filtering and preprocessing step was carried out. Then, the peptides were separated based on their MS1 response and labeled accordingly (Fig. 1A), further splitting it into training, validation, and test sets. Training and optimization were carried out selecting the best model. The concept behind the model’s architecture (Fig. 1B, S4) is to generate a more meaningful representation of the data through the encoder. The attention mechanism then identifies the relevance of each of the constituent elements of the input sequences. Finally, the new data representation and the attention weights are used by the decoder to make the predictions.

### 3.1. Base model performance on ProteomeTools data

The test set from the ProteomeTools data was used to do an initial evaluation of the base model, which achieved a categorical accuracy of 66%, based on the four classes established in this study. The base model appeared to be more prone to misclassifying weak flyers as non-flyers compared to intermediate or strong flyers (Fig. 1C inset). This suggests that the model is finding more similitudes between non-flyers and weak flyers. Moreover, the model had difficulties distinguishing between the different flyer classes, with 87% of the misclassified peptides falling into neighboring classes, especially for those near the threshold between classes (Fig. S5). The model had better performance classifying non-flyers (AUC = 0.97), followed by strong flyers (AUC = 0.90) and weak flyers (AUC = 0.85), while intermediate flyers (AUC = 0.78) were the most challenging (Fig. 1C). These results might be explained by the fact that the dataset was split based on MS1 intensity values, so the number of peptides in each class was balanced. As the intensity values are continuous (i.e. not ordinal data), the cut off points are blurry lines between classes. While threshold intensities were defined to build a model that could indicate the likelihood of peptide detectability and distinguish between weak, intermediate, and strong flyers, the data-dependent and non-absolute nature of this approach likely resulted in decreased classification performance.

Based on these challenges, the multi-class model output was transformed into a binary format by aggregating the probabilities of all flyer classes, without altering the architecture. The cumulative probability now represents the overall probability of a peptide being a flyer, while the probability of it being a non-flyer remains unchanged from its initial prediction. Although the final binary prediction is based on selecting the class with the highest probability; if any of the flyer classes (i.e., weak, intermediate, and strong flyers) holds the highest probability, the peptides are labeled as flyers; otherwise, they are categorized as non-flyers. After this modification, the model’s performance increased considerably having a binary accuracy of 94%, an MCC of 0.85, a precision of 96%, a recall of 97%, an F1 score of 96%, and an AUC of 0.97 (Fig. 1D). Thus, the model demonstrates greater effectiveness in distinguishing between flyers and non-flyers compared to distinguishing among the four different categorical labels. Converting the multi-class categorical prediction into binary format improved model performance, surpassing that of models trained directly on binary (flyer/non-flyer) labeled data (data not shown).

### 3.2. Inherent bias for synthesizability

Given that the base model was trained using data from a synthetic peptide library, a potential bias towards peptide synthesizability was investigated. This bias could lead to a tendency to predict peptides as flyers if they are easier to synthesize, while those that are more challenging to synthesize may be predicted more often as non-flyers. The ProteomeTools dataset was initially selected in part because of its homogeneous nature in terms of sample preparation, concentrations, and experimental setup and because it does not rely on additional factors such as digestion. This consistency allows, in principle, the elimination of any potential influence of peptide abundance in the learning process. In this context, biological datasets have the inconvenience of representing a wide range of protein concentrations, introducing peptide abundance as a variable factor that needs to be learned implicitly by the model. Moreover, the variability in site-specific protein digestibility introduces additional uncertainty into the ground truth of peptide presence and abundance.

Any bias towards synthesizability in the base model was explored using the predictions made on the test subset of the Sinitcyn *et al.* dataset (for details on the train/test split creation see Methods). For this, peptide synthesizability and selected/multiple reaction monitoring (SRM/MRM) experimental compatibility was compared with the correct and incorrect predictions made by the base model. 2,000 peptides were sampled for each possible outcome of the binary classification: correctly identified flyers or true positives (TPs), correctly predicted non-flyers or true negatives (TNs), incorrectly predicted flyers or false positives (FPs), and incorrectly predicted non-flyers or false negatives (FNs). To determine the synthesizability and SRM/MRM compatibility of peptides, ThermoFisher’s Peptide Synthesis and Proteotypic Peptide Analyzing Tool (online)^71^ was used. It is important to note that this tool does not support any peptide modifications and the quality of the analysis can be greatly impacted if the peptide sequence is longer than 25 AAs. Thus, the results are taken as an indication of peptide synthesizability and SRM/MRM compatibility rather than an absolute truth.

The experiment shows that TPs and FPs display very similar synthesizability and SRM/MRM compatibility profiles (Fig. 2A, 2C), although TPs have slightly better SRM/MRM compatibility. Thus, FPs represent peptides that should be well-suited for MS yet go undetected in experimental analysis. This discrepancy may not be directly attributed to the peptides themselves, but rather to factors such as peptide abundance, protein digestibility, sample complexity, and other variables related to the biological nature of the dataset. Additionally, the model may predict non-flyers as flyers due to their ease of synthesis and better suitability for MS analysis. While these two explanations are plausible, they are not mutually exclusive; it is possible that a combination of both contributes to this phenomenon. Both TNs and FNs tend to have poorer synthesizability and SRM/MRM compatibility than TPs and FPs, albeit TNs generally seems worse. Therefore, TNs might be predicted as such due to lower synthesizability and SRM/MRM compatibility. On the other hand, FNs tend to have worse synthesizability than TPs but are more compatible with MS than TNs. These observations suggest that peptide synthesizability might influence the predictive capabilities of the base model and indicate a potential inherent bias. Certain sequence elements may, however, affect peptide detectability through different mechanisms. For instance, the dimer motif asparagine-glycine (Asn-Gly) has been shown to not only reduce peptide synthesizability but also decrease peptide stability and may induce gas-phase reactions during ionization^55^. While all these mechanisms ultimately lead to a decreased MS response and, thus, a higher probability of peptides being predicted as non-flyers, differentiation between synthesizability and inherent peptide properties related to detectability is not possible with the base model. To address a potential bias towards synthesizability, the base model was fine-tuned by further training it on a subset of the Sinitcyn *et al.* dataset (for details on the train/test split creation see Methods). Upon testing the fine-tuned model and closely examining the results, it became apparent that there was a noticeable reduction in the bias towards synthesizability. This was especially evident in the more evenly distributed synthesizability and SRM/MRM compatibility observed among TNs and FNs when compared with the base model results (Fig. 2A - 2D). However, by looking at the distribution of TPs and FPs, there was minimal variation in the results between the two models, suggesting that a potential bias towards synthesizability may still persist. Nonetheless, this consistency in the results also leads to the hypothesis that there could be an inherent correlation between synthesizability and detectability.

**Figure 2.**
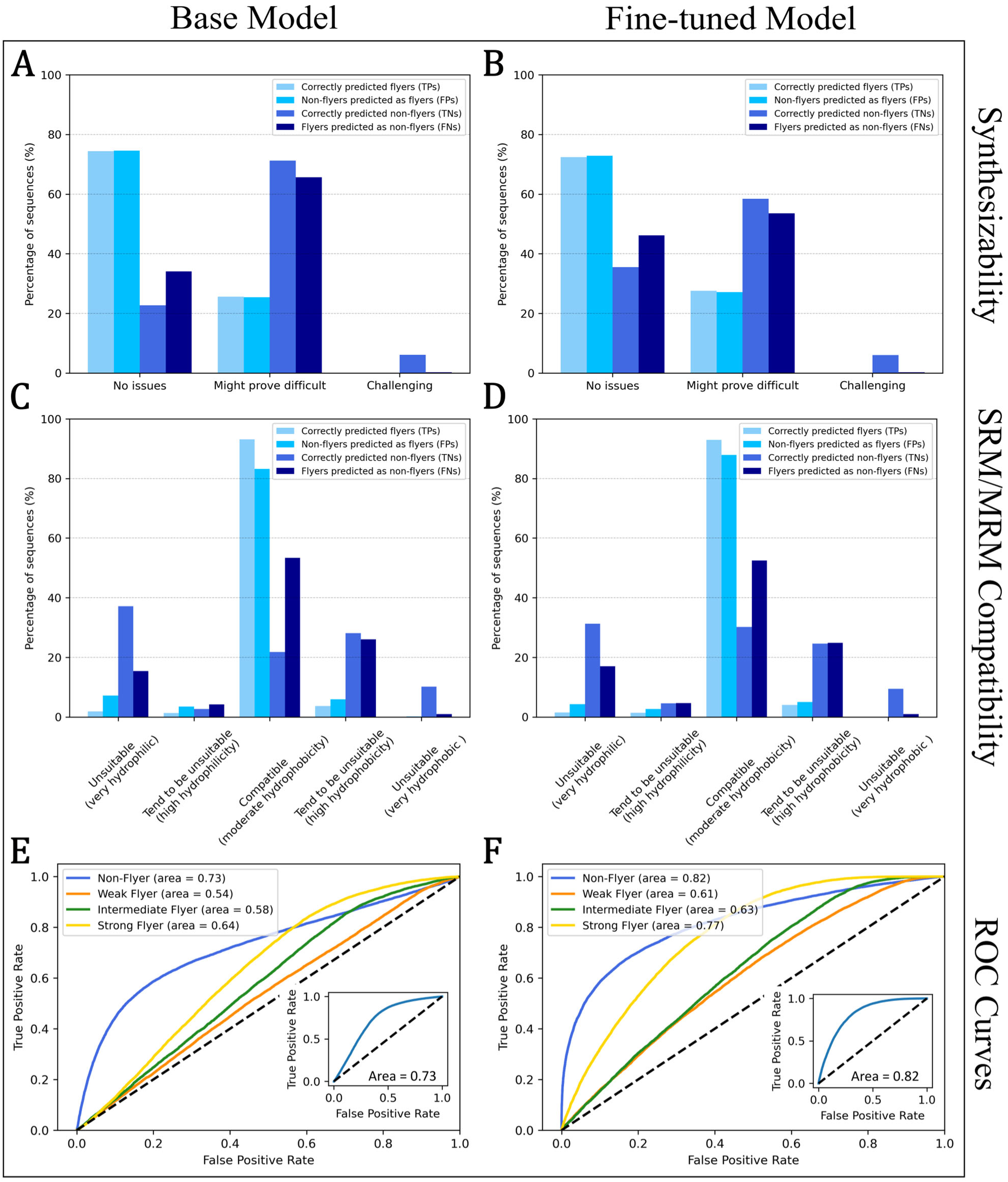
Comparative performance of base and fine-tuned “Pfly” model. Performance of the base model (left column) and the fine-tuned “Pfly” model (right column) based on predicted synthesizability (top row), SRM/MRM compatibility (middle row) and ROC curves for multi-class and binary predictions (bottom row), evaluated on the Sinitcyn *et al.* test dataset., Synthesizability for correctly and incorrectly predicted flyers and non-flyers by the base model (A) and the fine-tuned model (B). SRM/MRM compatibility prediction for correctly and incorrectly predicted flyers and non-flyers by the base model (C) and the fine-tuned model (D). ROC curves for multi-class and binary (insets) predictions obtained from the base model (E) and the fine-tuned model (F) for non-flyers (blue), weak flyers (orange), intermediate flyers (green) and strong flyers (yellow) along with the area under the curve for the predictions of the individual classes. For panels A-D, the data is color-coded to represent correctly predicted flyers (true positives, TPs), non-flyers incorrectly predicted as flyers (false positives, FPs), correctly predicted non-flyers (true negatives, TNs), and flyers incorrectly predicted as non-flyers (false negatives, FNs), with each category representing a random sample of 2,000 peptides.

Another aspect worth considering is the impact of the fine-tuning process itself. By employing a biological and non-equimolar dataset for fine-tuning, there is a possibility that biases related to protein concentrations and digestibility, and therefore also peptide abundance were inadvertently introduced. Thus, peptides containing the Asn-Gly dimer motif within this subsample of 8,000 peptides were studied, allowing a deeper look at the relationship between synthesizability, detectability, and abundance. In this specific example, a reduction in the effect of synthesizability in the fine-tuned model can be observed, with a substantial decrease in FNs (from 50% to ∼20%) and an increase in TPs (from 10% to ∼40%) compared to the base model (Fig. S6A). Although it is not possible to prove a relationship between synthesizability and detectability, it is evident that non-flyers more frequently originate from proteins with lower intensity responses, as shown by the log-intensity distributions of TNs and FPs for both the base model and the fine-tuned model (Fig. S6B). Overall, there is a noticeable improvement in the performance of the fine-tuned model over the base model on the Sinitcyn *et al.* test dataset (Fig. 2E - 2F). The area under the ROC curve increases from 9% (intermediate flyers, 0.58 to 0.63) to 20% (strong flyers, 0.64 to 0.77) for the multi-class predictions and 12% (0.73 to 0.82) for the binary predictions after fine-tuning. This shows that the fine-tuning improved the model’s predictive capabilities on biological data.

### 3.3. Model benchmarking

The predictive capability of the fine-tuned model (Pfly) was benchmarked against the base model and several other state-of-the-art peptide detectability models, namely DeepMSPeptide^48^, PepFormer^23^, and DeepDetect^72^. The tools used to evaluate performance were implemented according to the algorithmic specifications provided in their source code. Initially, the Sinitcyn *et al.* test dataset was used to compare the results of the different tools. Pfly reached an accuracy, MCC, and AUC of 0.78, 0.50, and 0.82, respectively, outperforming the other tools with accuracy values ranging from 0.67 to 0.72, MCC values from 0.24 to 0.34, and AUC values from 0.64 to 0.73, among other metrics assessed (Fig. 3A, Table S1). Since Pfly was fine-tuned with a partition of the Sinitcyn *et al.* dataset, the next sensible step was to test all the models with a more independent dataset for which the test set of the Wang *et al.* dataset^58^ (see Methods) was used. On this dataset, Pfly achieved an accuracy of 0.68, an MCC of 0.43, and an AUC of 0.78, outperforming the other tools with values ranging between 0.57 to 0.64 for accuracy, 0.23 to 0.31 for MCC, and 0.64 to 0.73 for AUC, among other evaluated metrics (Fig. 3B, Table S2). Overall, Pfly showed a notably higher predictive capacity across both datasets in comparison with the other tools used in this study, including the base model. Thus, fine-tuning had a positive effect on our model, improving its performance not only on the dataset from which the fine-tuning subset originates, but also on an independent dataset that the model had never seen during training.

**Figure 3.**
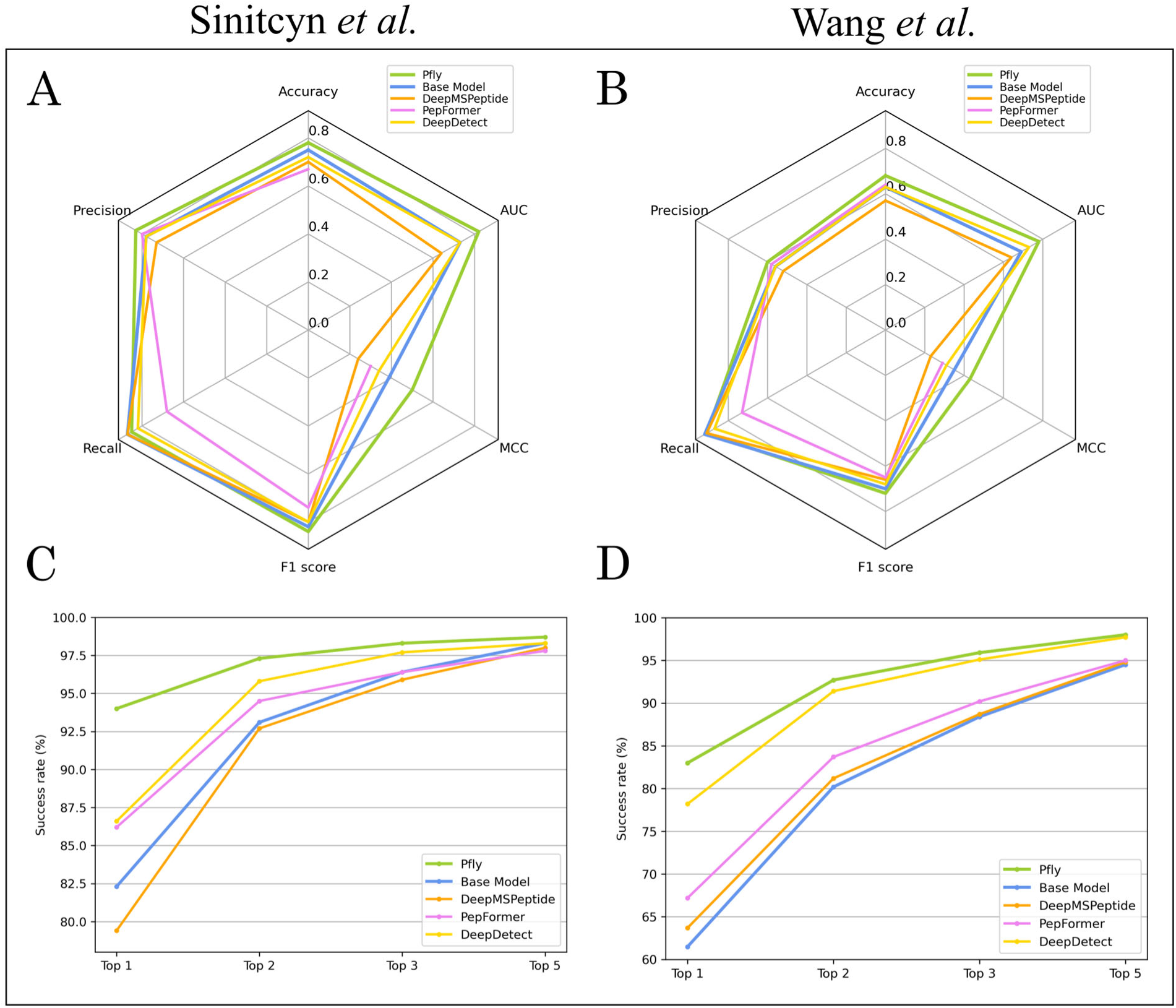
Performance comparison of the base and Pfly models against other tools. (A) Comparison on the Sinitcyn *et al.* test dataset and (B) on the Wang *et al*. test dataset using accuracy, precision, recall, F1 score, Matthews Correlation Coefficient (MCC), and area under the curve (AUC) as evaluation metrics for the individual models. (C) Comparison on the Sinitcyn *et al.* test dataset and (D) on the Wang *et al.* test dataset showing the Top 1, Top 2, Top 3, and Top 5 ranked peptides for the individual models. The “Top X” peptides refers to the certainty of experimentally identifying a peptide within the X top ranked peptides predicted as flyers by the individual models.

To take a more practical approach on the applicability of Pfly, its performance was measured based on the top ranked peptides predicted as flyers per protein. The “Top X” metric refers to the X number of peptides from a specific protein with the highest predicted probability of being a flyer according to a given model. If one or more of these top X peptides are identified experimentally, the model is considered to have correctly predicted a flyer for that particular protein. Therefore, the Top X metric provides a percentage of certainty or success rate, indicating how effectively a model can identify a flyer within the top X predicted peptides for a specific protein, with the peptides from each protein determined through an *in silico* digest. This approach provides an evaluation of a model’s ability to accurately predict peptides suitable for e.g. targeted analysis in MS-based proteomics. Similarly to previous analyses, the test splits from the Sinitcyn *et al.* and Wang *et al.* datasets were used for benchmarking. Based on the Top X metric, Pfly outperformed all the other models across both datasets (Fig. 3C-D). The most noticeable difference was observed with the Top 1 metric, where Pfly scored 94.0% on the Sinitcyn *et al.* dataset and 83.0% on the Wang *et al.* dataset. In comparison the other tools ranged from 79.4% to 86.6% on the Sinitcyn *et al.* dataset, and from 63.7% to 78.2% on the Wang *et al.* dataset (Table S3, S4). With the Top X metric, the base model exhibited comparatively weaker performance compared to the other tools, having DeepDetect as a runner-up. This contrasts with the performance of the base model when evaluating using other metrics (Fig. 3A-B).

### 3.4. Effect of protein abundance on the model’s performance

The improved performance of the model after fine-tuning (Fig. 3) could indicate that the base model indeed had a bias towards peptide synthesizability. However, it is also essential to consider how the abundance of proteins and peptides may impact the results. To do so, the performance of Pfly was evaluated on the two test datasets used in the previous analyses by examining the precision per protein. Precision per protein is defined as the ratio between the number of correctly predicted flyers (TPs) and the total number of predicted flyers (TPs + FPs) for individual proteins. This assessment assumes that higher protein-level abundance will increase the abundance of PTPs, and that greater peptide abundance enhances the likelihood of experimentally identifying flyer peptides. As such, precision per protein allows for the evaluation of abundance-related factors, based on the hypothesis that precision per protein will increase with higher protein abundance. iBAQ values were used as a measure of protein-level abundance, as they are considered a good proxy of molar abundance within a particular dataset. The analysis was conducted by comparing the peptides per protein identified across all experiments with those identified in a single experiment. For the test split from the Sinitcyn *et al.* dataset, the flyers identified across all six cell lines were compared with those identified in one randomly selected cell line (GM12878) digested with trypsin. In the same fashion, for the Wang *et al.* dataset, the flyers identified across all tissues were compared with those identified in the randomly selected tonsil tissue. This specific cell line and tissue were selected to exemplify if there was any correlation between protein abundance and peptide detectability.

In the Sinitcyn *et al.* dataset, there is a clear difference in precision per protein when using data from a single cell line compared to combining information from all cell lines (Fig. 4A-B). Utilizing data from all cell lines boosts the likelihood of identification of the constituent peptides of each protein. Furthermore, a positive correlation between protein abundance (iBAQ values) and precision per protein can be seen (Fig. 4A). This correlation is also evident in the Wang *et al.* dataset (Fig. 4C). These observations suggest that the influence of protein abundance is more pronounced when analyzing data from a single experiment. Here, the abundance of individual proteins is restricted within a specific and narrow range, compared to a larger dataset from different cell lines/tissues, where protein abundance will cover a wider dynamic range and individual proteins are more likely to be abundant in at least one subset. This is evident from the increase in both iBAQ values (right-shift), and precision (up-shift) observed when moving from a single cell line/tissue analysis (Fig. 4A, C) to all cell lines/tissues (Fig. 4B, D). In the single sample analyses, precision is hampered by low-abundance peptides misclassified as FPs. These FPs, which were actually TPs in all cell line/tissue experiments, highlight the model’s capacity to correctly predict flyer peptides that might not be detected for reasons unrelated to their ability to fly, such as protein concentration or digestibility—a trend observed across both the Sinitcyn *et al.* and Wang *et al.* datasets (Fig. 4C, D). In contrast, a decreasing tendency in recall was observed at higher protein abundances (Fig. S7), indicating an increase in FNs. This suggests that peptides that the model would expect to be non-flyers at higher abundance are indeed detectable, further accentuating the relevance of abundance in the model’s results.

**Figure 4.**
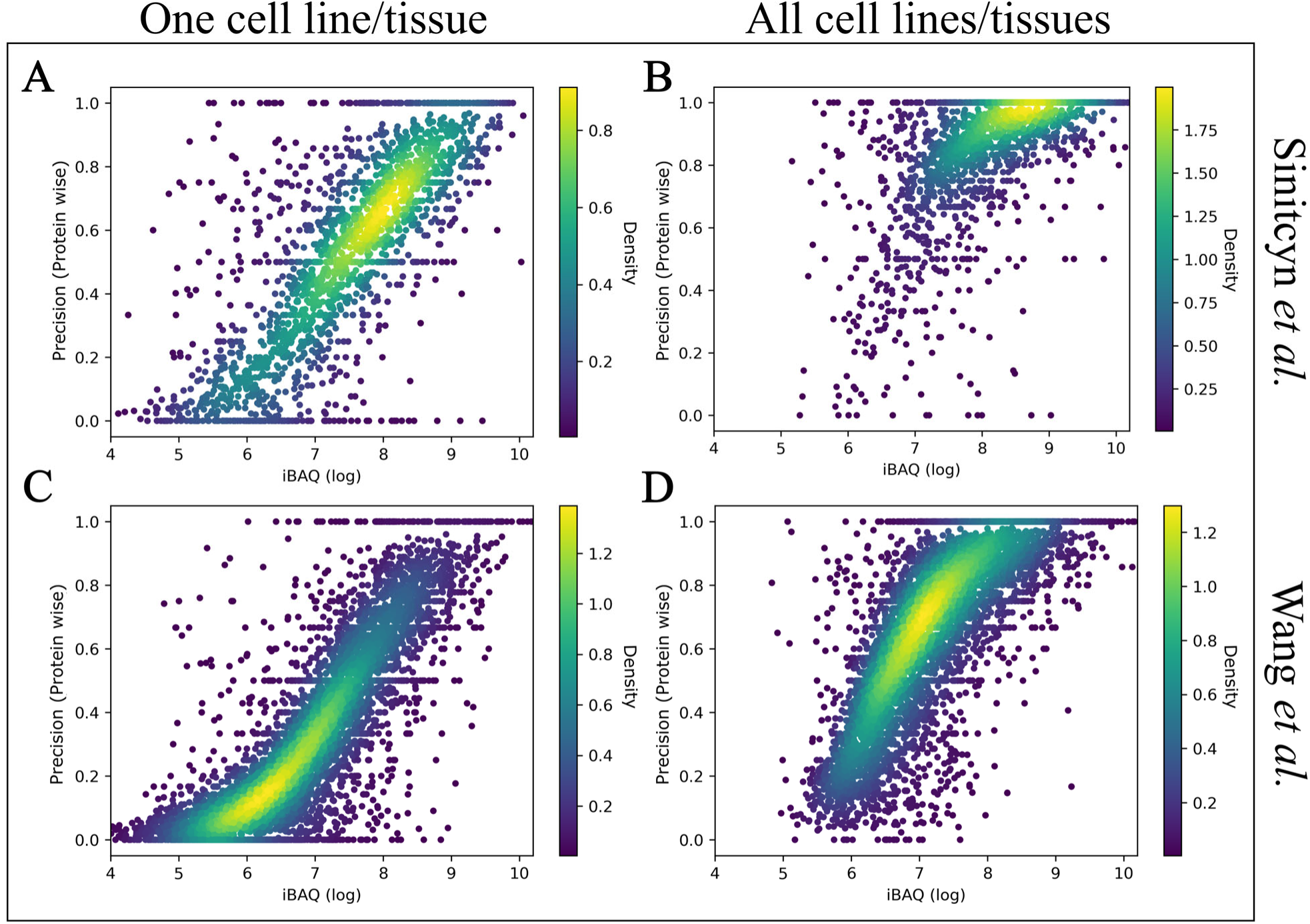
Precision per protein versus the corresponding protein iBAQ values. Precision per protein is calculated as the ratio of correctly predicted flyers (TPs) to the total number of predicted flyers (TPs + FPs) for specific proteins. This metric is obtained by evaluating the Sinitcyn *et al.* test dataset using labels from the GM12878 cell line (A) and from all six cell lines (B), as well as the Wang *et al.* test dataset using labels from the tonsil tissue (C) and from all thirty tissues (D). Precision per protein is plotted against the corresponding protein iBAQ values. Each dot represents an individual protein (determined by its corresponding peptides), and the data distribution is color-coded by point density from low (dark blue) to high (yellow).

### 3.5. Effect of MS raw data processing on peptide identification

In MS analysis, one factor that influences missed peptide identification is the processing of raw data. Common database search engines for bottom-up proteomics evaluate scores based on peptide fragments, fitting experimental spectra against *in silico* generated spectra to determine matches^73^. However, these scores can be suboptimal, leading to the loss of correct matches. To address this, machine learning techniques have been used to develop models for rescoring, enhancing rates of peptide and protein identification^70,74^.

To explore this possibility and investigate the effect of peptide scoring (using the Andromeda^75^ search engine integrated in MaxQuant^76^), Oktoberfest^70^ was utilized to rescore the Sinitcyn *et al.* test dataset to assess how many FPs predicted by Pfly, were actually TPs missed during MS raw data processing.

From the 2,000 proteins used for testing, 60,185 peptides were obtained from an *in silico* tryptic digest whereof 19,339 peptides (32%) were not identified in the dataset and consequently labeled as non-flyers. Of the labelled non-flyers, 3,120 peptides (16%) were identified by rescoring whereof 1,461 (47%) were predicted as flyers by the fine-tuned (Pfly) model (Table S5). Upon recalculating all previous metrics, a discernible enhancement in performance was observed across most metrics. However, recall showed a notable decrease, negatively impacted by the increment of FNs. However, minimal differences were noted in accuracy and F1 score (Fig. 5A, Table S6). This improvement underscores the efficacy of the rescoring process in rectifying misclassifications and enhancing overall model performance. Considering the Top X peptides metrics, a slight but consistent improvement in all the Top X peptides was observed (Fig. 5B, Table S7). These observations serve as a clear indicator of the multitude of factors influencing peptide detectability in MS analysis, including protein abundance and the processing of raw MS data. Consequently, these factors are likely to have a significant impact on the outcomes of predictive models like Pfly. Understanding and addressing these influencing factors is crucial towards refining and optimizing the performance of such models. In navigating this complexity, Pfly emerges as a standout solution, leveraging a robust baseline model that can be fine-tuned with external data from diverse samples, acquisition conditions, and search engines. This adaptability enhances Pfly’s versatility and predictive accuracy across various experimental settings, enabling it to continuously evolve and maintain its reliability as a tool for advancing proteomic research and applications, ensuring its relevance and effectiveness in cutting-edge proteomics.

**Figure 5.**
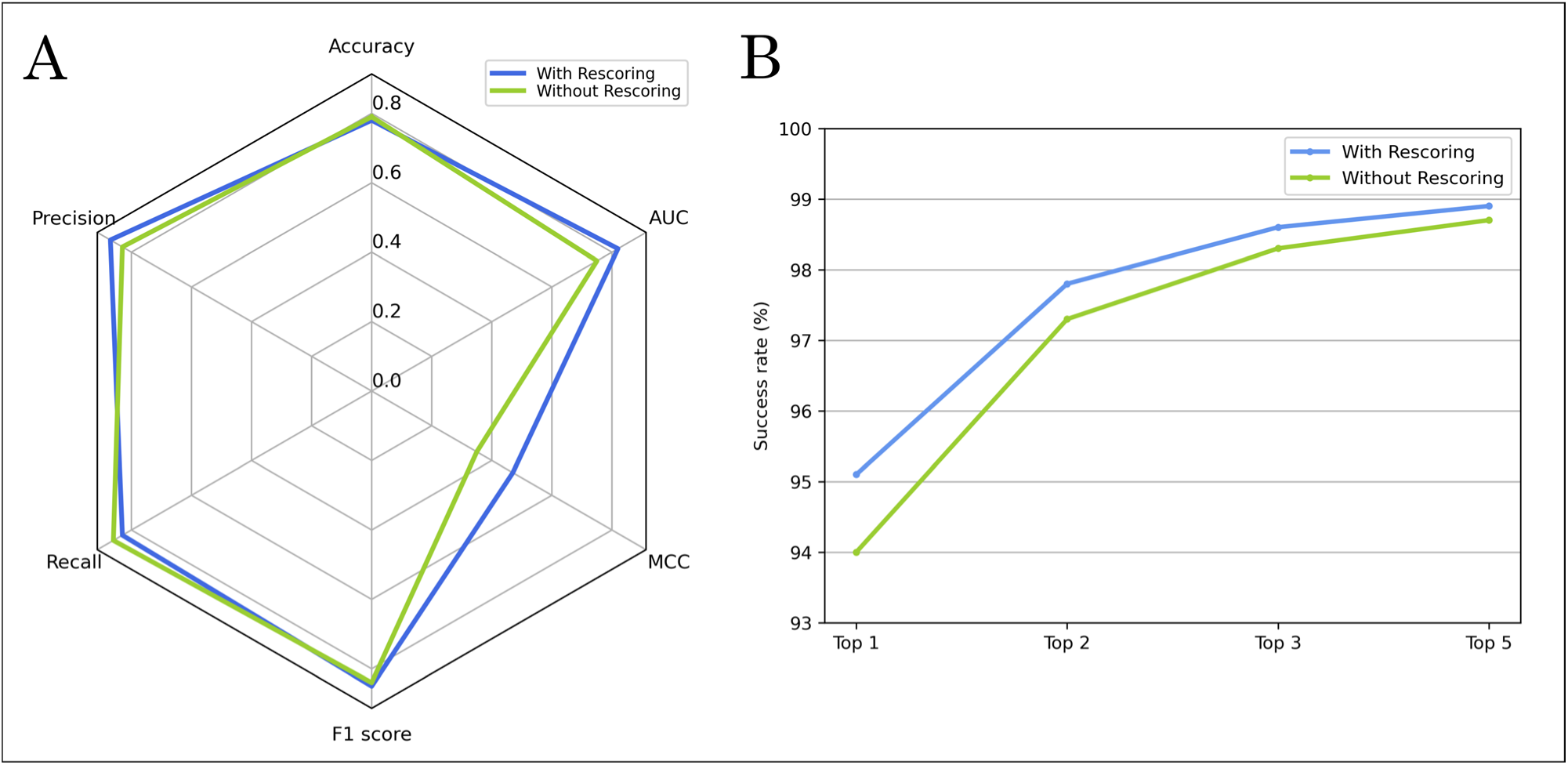
Performance comparison of Pfly on the rescored and initial Sinitcyn *et al.* test datasets. (A) Performance is shown using accuracy, precision, recall, F1 score, Matthews Correlation Coefficient (MCC), and area under the curve (AUC) for both the original and rescored labels. (B) Performance is shown for Top 1, Top 2, Top 3, Top 5 peptides with both the original and rescored labels. Top X peptides refers to the certainty of finding a flyer among the X highest ranked peptides predicted by a model.

## 4. Conclusions

In this study, we present Pfly, a model for peptide detectability prediction based on an encoder-decoder architecture with an attention mechanism. Pfly accurately predicts whether a peptide is a flyer or a non-flyer while providing the probability assigned to the different classes defined in this study (non-, weak, intermediate, and strong flyer). A base model was initially built using a dataset generated from a synthetic peptide library intended to be equimolar. However, it was found to have a bias towards peptide synthesizability. Therefore, the model was fine-tuned using a biological dataset, which improved its performance and resulted in the final model, Pfly. In a comparative evaluation with other tools, Pfly showed improved performance across the vast majority of the evaluation metrics, establishing its value as a reliable tool for predicting peptide detectability in MS analysis. Moreover, we establish that there are other factors influencing the performance of our model, such as protein abundance. Peptides originating from low-abundance proteins may be misclassified not due to their ability to fly, but rather because of their low concentration in the sample. Additionally, we explored how the suboptimal performance of search engines can detrimentally impact MS analysis by leading to missed peptide identifications, which can, in turn, affect protein identification and quantification. Peptide detectability is influenced by the entire pipeline, encompassing factors from sample characteristics such as abundance, to the specific database search engine utilized, with each intermediate step playing a crucial role. Thus, developing a generic model capable of accounting for all these complexities poses challenges, necessitating methods that facilitate the fine-tuning of models to specific experimental conditions. These complexities also underscores the uncertainty in results due to the various intermediate steps, which can inadvertently bias outcomes when models are trained and tested on similar datasets. For instance, peptides currently predicted as undetectable may be the result of a model being overfitted to a specific dataset, leading to biases that impair its generalizability to other datasets. Consequently, further research is imperative to thoroughly investigate the cumulative effects of each step in the analytical process. This highlights the non-trivial nature of evaluating models based on comprehensive properties like detectability, which can unintentionally be influenced by tool selection, equipment, and experimental design. Emphasizing the necessity for ongoing improvement in protocols, MS technology and data processing tools, these insights underscore the importance of aligning advancements in analytical methods with the evolving complexities of peptide detection and quantification.

Pfly represents a significant advancement in the field of computational proteomics. Integrated into a comprehensive pipeline, it facilitates the implementation of the model, encompassing training, optimization, and customization. This flexibility allows users to fine-tune models according to specific experimental conditions, thereby generating a lab-specific versions with superior performance. Researchers can use this tool to improve protein identification in MS experiments by distinguishing true peptide signals from noise and supporting quality control by assessing peptide detectability and removing unreliable data. It can also play a crucial role in biomarker discovery by predicting consistently detected peptides, thus helping to identify potential biomarkers relevant for diagnosing diseases or monitoring treatment responses. In comparative proteomics studies, Pfly can assess protein differential expression, providing insights into biological responses and experimental interventions. It can also optimize experimental designs in targeted proteomics experiments by accurately identifying detectable peptides. Furthermore, it can assist in interpreting large-scale proteomic datasets, prioritizing peptides and proteins, for instance, to provide a deeper understanding of molecular mechanisms and pathways. Pfly’s capabilities can extend to food science for quality control and nutritional analysis, as well as to other fields such as pharmaceuticals, agriculture, and environmental sciences, where precise protein and peptide identification is essential. These diverse applications underscore the broad impact and significance of Pfly in advancing proteomics, biological science, and numerous related disciplines.

## 5. Author contributions

**Naim Abdul-Khalek:** Conceptualization, Methodology, Software, Validation, Formal analysis, Investigation, Data Curation, Writing - Original Draft, Writing - Review & Editing, Visualization. **Mario Picciani:** Formal analysis, Investigation, Data Curation, Writing - Review & Editing. **Reinhard Wimmer:** Conceptualization, Resources, Writing - Review & Editing, Supervision. **Michael Toft Overgaard:** Conceptualization, Resources, Writing - Review & Editing, Supervision, Project administration, Funding acquisition. **Mathias Wilhelm:** Conceptualization, Methodology, Resources, Writing - Review & Editing, Supervision. **Simon Gregersen Echers:** Conceptualization, Methodology, Resources, Writing - Review & Editing, Supervision, Project administration, Funding acquisition.

## 6. Data availability

The data used in this study was acquire from the ProteomeXchange Consortium with the identifiers PXD004732, PXD010595, PXD021013, PXD024364, and PXD010154 and the supplementary material from corresponding publications^52–54^.

## 7. Code availability

Source code and scripts are available on GitHub at https://github.com/wilhelm-lab/dlomix.

## 9. Acknowledgements

This work was in part funded by the Karl Pedersen & Hustrus Industrifond (grant number DI-2019– 07020), the European Research Council (ERC; grant number 101077037), the German Federal Ministry of Education and Research (BMBF; grant number 031L0305A), and the Elite Network of Bavaria (ENB; grant number F-6-M5613.6.K-NW-2021-411/1/1)

