## Supplementary information for "To fly, or not to fly, that is the question: A deep learning model for peptide detectability prediction in mass spectrometry"

### For

**Figure S1.** Flow chart of the data filtering process for the ProteomeTools dataset.
**Figure S2.** Flow chart of the data filtering process for the Sinitcyn et al. dataset.
**Figure S3.** Flow chart of the data filtering process for the Wang et al. dataset.
**Figure S4.** General Scheme of Pfly Neural Network Architecture.
**Figure S5.** Boxplot showing intensity distribution of correctly classified and misclassified peptides.
**Figure S6.** Comparative performance of the base and fine-tuned (Pfly) models of peptides sequences containing the "NG" dimer motif.
**Figure S7**. Recall per protein versus the corresponding protein iBAQ values.

**Table S1**. Performance metrics of base and fine-tuned (Pfly) models and other tools used for benchmarking on the Sinitcyn *et al.* dataset.
**Table S2**. Performance metrics of base and fine-tuned (Pfly) models and other tools used for benchmarking on the Wang *et al.* dataset.
**Table S3**. Top flyers accurately predicted (%) by the base and fine-tuned (Pfly) models and the other tools used for benchmarking on the Sinitcyn *et al.* dataset.
**Table S4**. Top flyers accurately predicted (%) by the base and fine-tuned (Pfly) models and the other tools used for benchmarking on the Wang *et al.* dataset.
**Table S5**. Partitions of the Sinitcyn *et al.* test dataset.
**Table S6**. Performance metrics of the fine-tuned (Pfly) model on the originally labeled and rescored Sinitcyn *et al.* test dataset.
**Table S7**. Top flyers accurately predicted (%) by the fine-tuned (Pfly) model on the originally labeled and rescored Sinitcyn *et al.* test dataset.

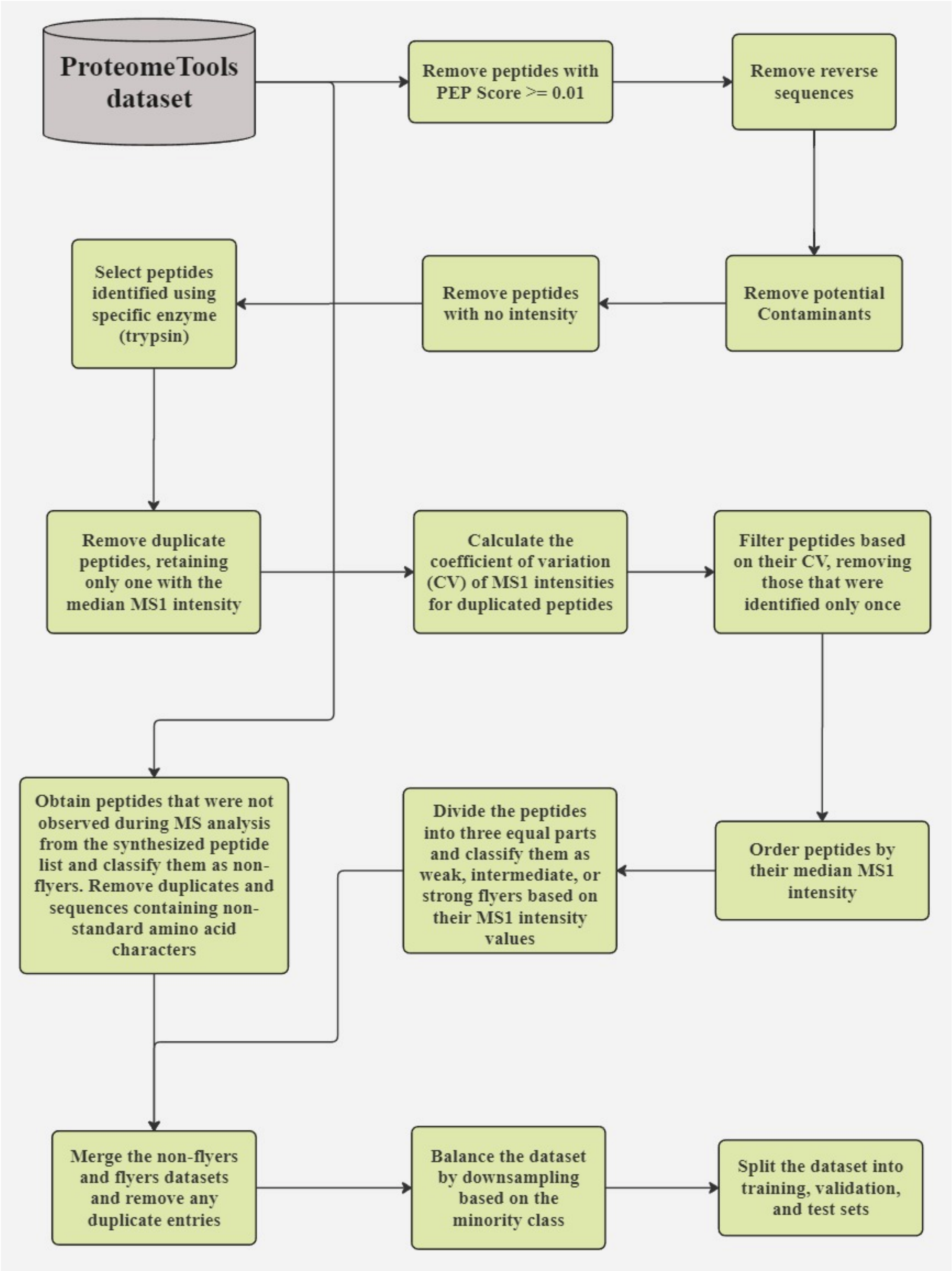


**Figure S5.** Flow chart of the data filtering process for the ProteomeTools dataset[1–3].

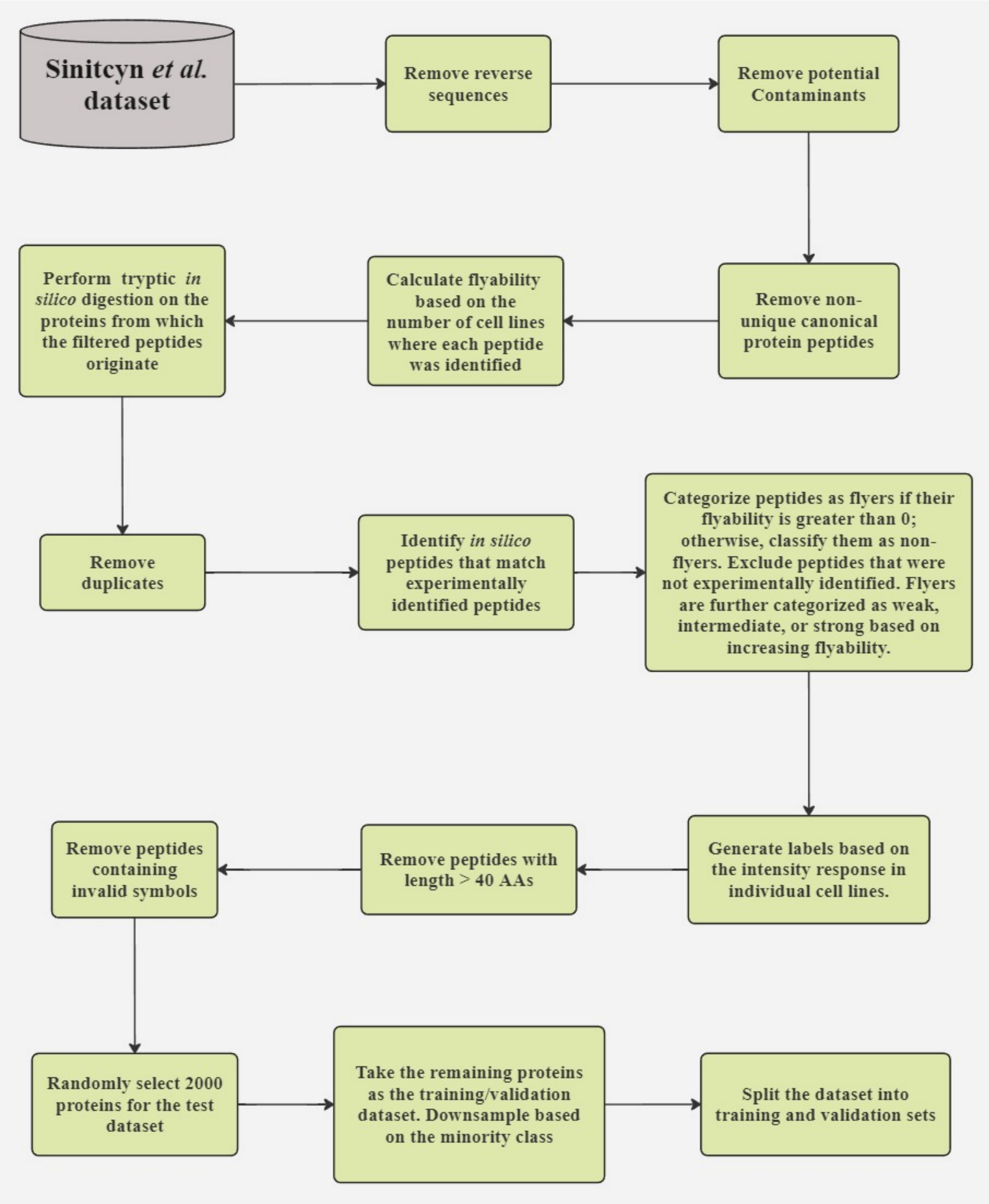


**Figure S6.** Flow chart of the data filtering process for the Sinitcyn *et al.* dataset[4].

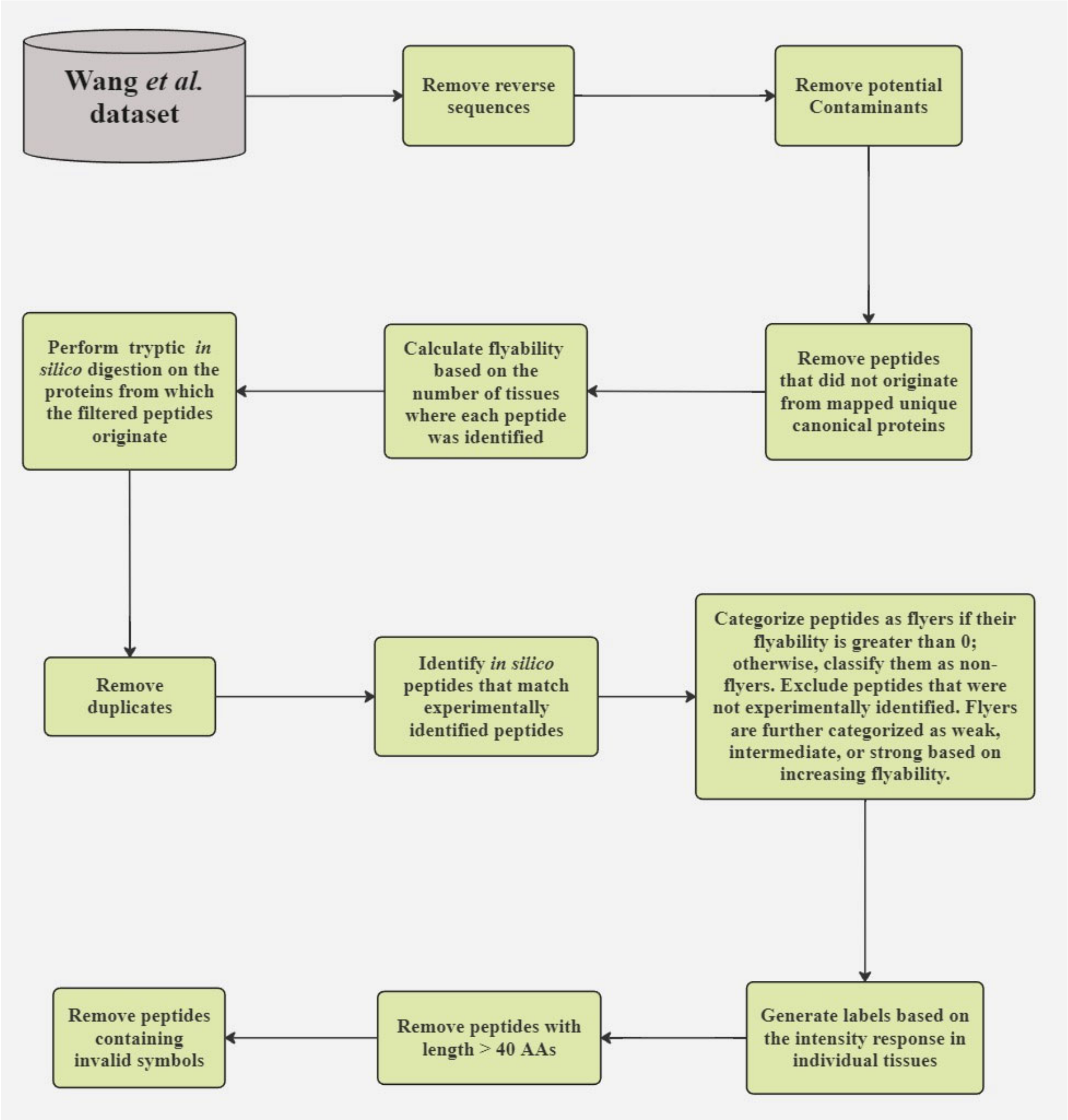


**Figure S7.** Flow chart of the data filtering process for the Wang *et al.* dataset[5].

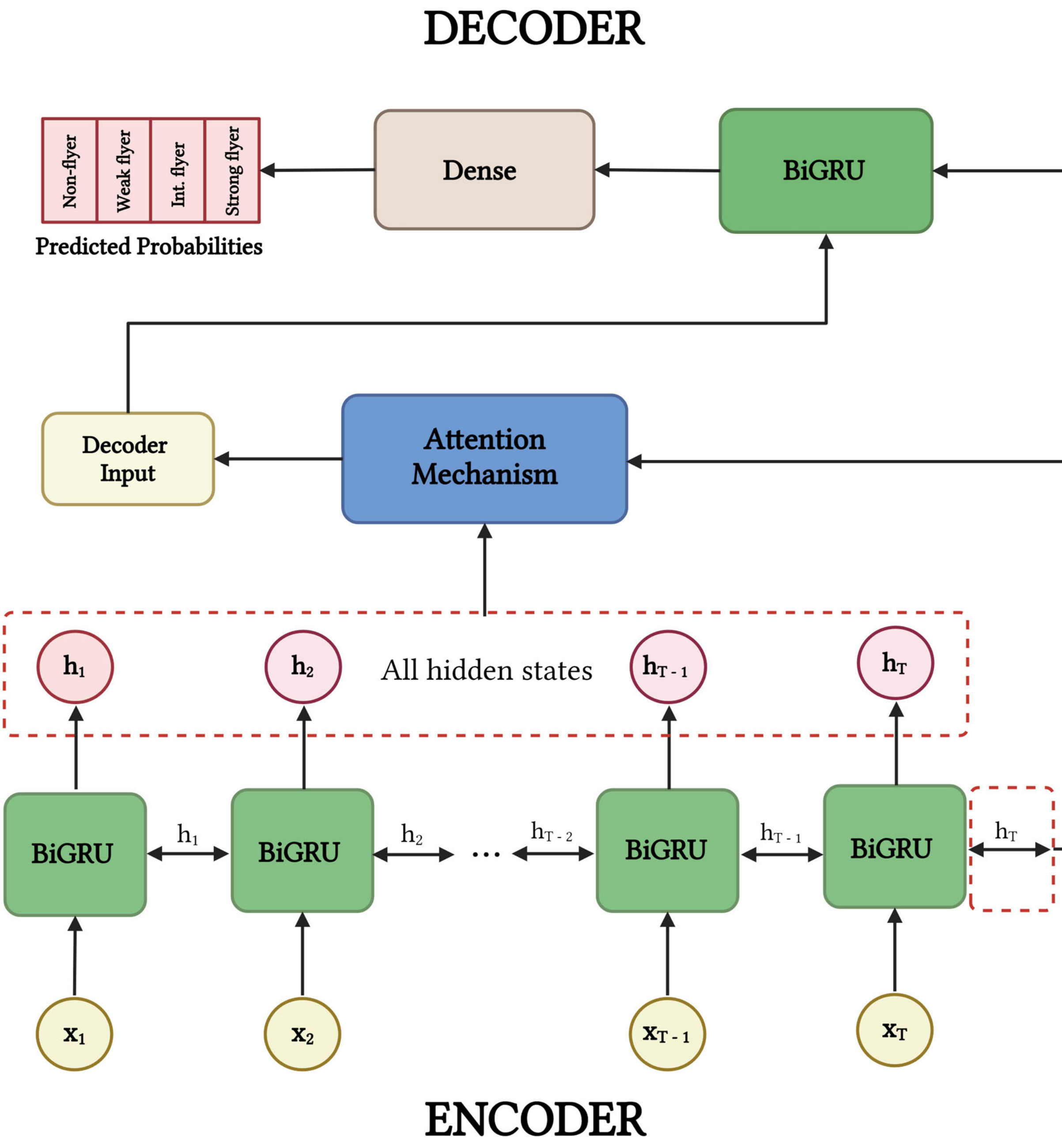


**Figure S8.** General Scheme of Pfly Neural Network Architecture adapted from Abdul-Khalek *et al.*[6]. The Pfly model utilizes an encoder-decoder framework with an attention mechanism. The encoder comprises a single bidirectional GRU layer (BiGRU) that processes the inputs ($x_1$, ..., $x_T$) and generates the corresponding hidden states ($h_1$, ..., $h_T$). The initial state of the decoder is set to the last hidden state of the encoder. This, along with all encoder hidden states, is used by the attention mechanism to compute a context vector that serves as input for the decoder. The decoder consists of one BiGRU layer and a dense layer with a softmax activation function. This dense layer outputs a vector of four probabilities, each representing the likelihood of one of the four classes: non-flyer, weak flyer, intermediate flyer, and strong flyer. The decoder performs only one iteration since its output is not a sequence.

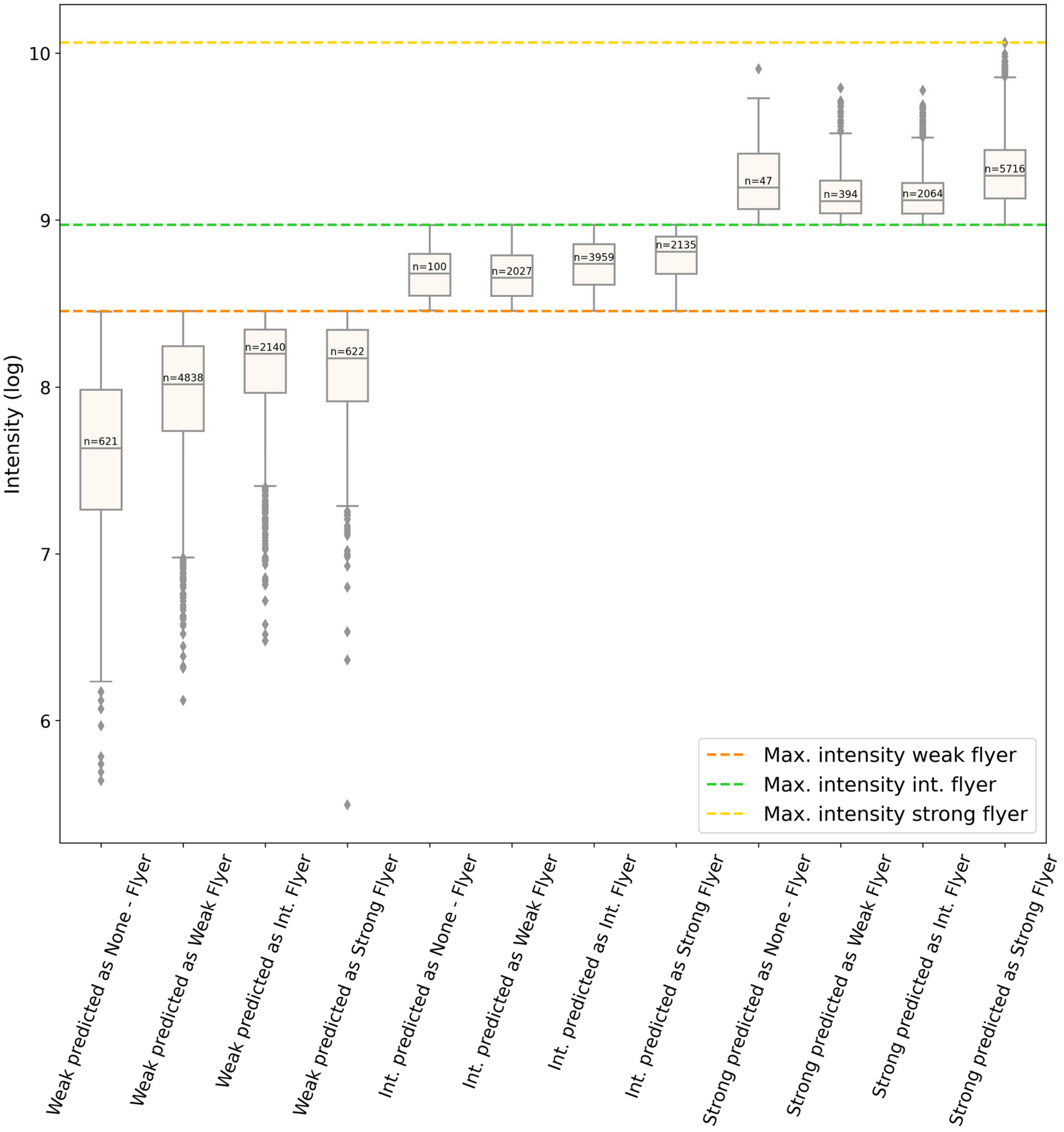


**Figure S5.** Boxplot showing intensity distribution of correctly classified and misclassified peptides from the test dataset from ProteomeTools[1–3].

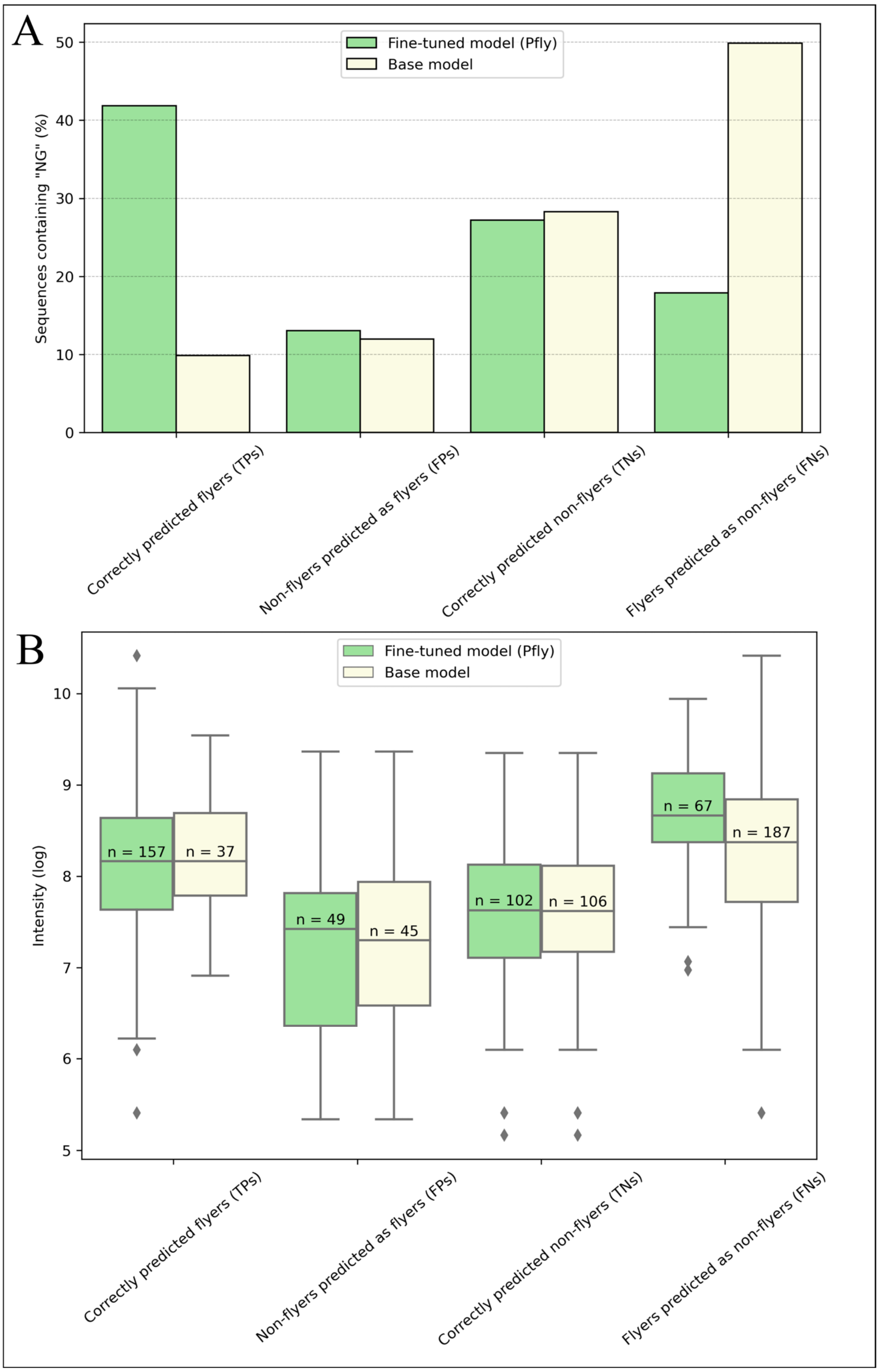


**Figure S6.** Comparative performance of the base and fine-tuned (Pfly) models of peptides sequences containing the "NG" dimer motif. (A) Correctly and incorrectly classified flyers and non-flyers by the base and fine-tuned model. (B) Protein log-transformed intensity profile for correctly and incorrectly classified flyers and non-flyers by the base and fine-tuned models. The results presented were obtained by evaluating both the base and fine-tuned models using a randomly selected subset of 8,000 peptides from the Sinitcyn *et al.* test dataset[4], focusing exclusively on peptide sequences that contain the "NG" dimer motif.

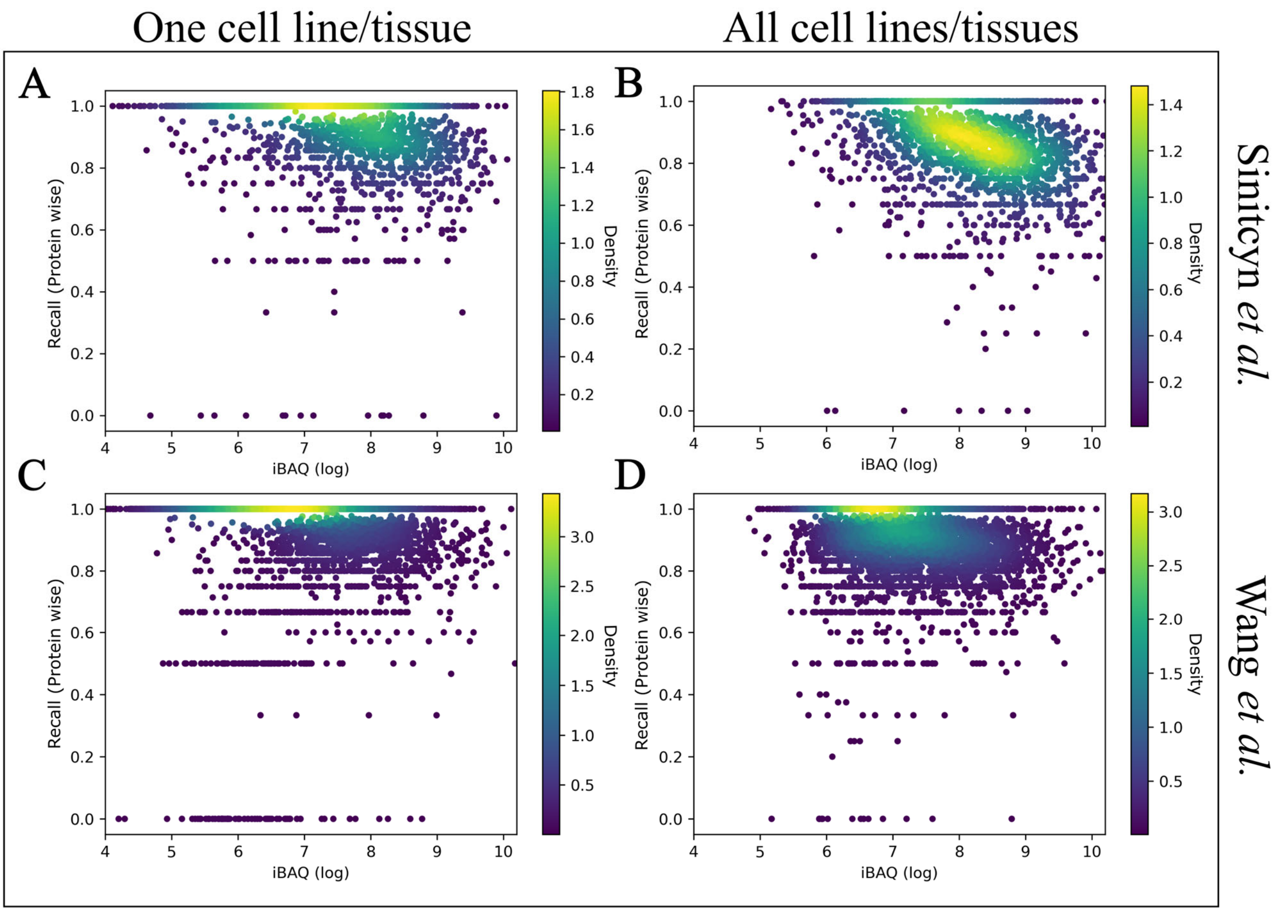


**Figure S7. Recall per protein versus the corresponding protein iBAQ values.** Recall per protein is calculated as the ratio of correctly predicted flyers (TPs) to the total number of actual flyers (TPs + FNs) for specific proteins. This metric is obtained by evaluating the Sinitcyn *et al.* test dataset using labels from the GM12878 cell line (A) and from all six cell lines (B), as well as the Wang *et al.* test dataset using labels from the tonsil tissue (C) and from all thirty tissues (D). Precision per protein is plotted against the corresponding protein iBAQ values. Each dot represents an individual protein (determined by its corresponding peptides), and the data distribution is color-coded by point density from low (dark blue) to high (yellow).

**Table S1**. Performance metrics of base and fine-tuned (Pfly) models and other tools used for benchmarking on the Sinitcyn *et al.* dataset[4], including accuracy, precision, recall, F1 score, Matthews Correlation Coefficient (MCC), and area under the curve (AUC).

| Model | Accuracy | Precision | Recall | F1 score | MCC | AUC |
|---|---|---|---|---|---|---|
| Base Model | 0.75 | 0.78 | 0.87 | 0.82 | 0.39 | 0.73 |
| Pfly | 0.78 | 0.83 | 0.85 | 0.84 | 0.50 | 0.82 |
| DeepMSPeptide | 0.70 | 0.73 | 0.87 | 0.80 | 0.24 | 0.64 |
| PepFormer | 0.67 | 0.80 | 0.68 | 0.74 | 0.30 | - |
| DeepDetect | 0.72 | 0.78 | 0.82 | 0.80 | 0.34 | 0.73 |

**Table S2**. Performance metrics of base and fine-tuned (Pfly) models and other tools used for benchmarking on the Wang *et al.* dataset[5], including accuracy, precision, recall, F1 score, Matthews Correlation Coefficient (MCC), and area under the curve (AUC).

| Model | Accuracy | Precision | Recall | F1 score | MCC | AUC |
|---|---|---|---|---|---|---|
| Base Model | 0.63 | 0.56 | 0.92 | 0.70 | 0.35 | 0.69 |
| Pfly | 0.68 | 0.60 | 0.91 | 0.72 | 0.43 | 0.78 |
| DeepMSPeptide | 0.57 | 0.52 | 0.91 | 0.66 | 0.23 | 0.64 |
| PepFormer | 0.64 | 0.58 | 0.73 | 0.65 | 0.29 | - |
| DeepDetect | 0.63 | 0.56 | 0.87 | 0.68 | 0.31 | 0.73 |

**Table S3**. Top flyers accurately predicted (%) by the base and fine-tuned (Pfly) models and the other tools used for benchmarking on the Sinitcyn *et al.* dataset[4].

| Model | Top 1 (%) | Top 2 (%) | Top 3 (%) | Top 5 (%) |
|---|---|---|---|---|
| Base Model | 82.3 | 93.1 | 96.4 | 98.3 |
| Pfly | 94.0 | 97.3 | 98.3 | 98.7 |
| DeepMSPeptide | 79.4 | 92.7 | 95.9 | 98.0 |
| PepFormer | 86.2 | 94.5 | 96.4 | 97.8 |
| DeepDetect | 86.6 | 95.8 | 97.7 | 98.3 |

**Table S4**. Top flyers accurately predicted (%) by the base and fine-tuned (Pfly) models and the other tools used for benchmarking on the Wang *et al.* dataset[5].

| Model | Top 1 (%) | Top 2 (%) | Top 3 (%) | Top 5 (%) |
|---|---|---|---|---|
| Base Model | 61.5 | 80.2 | 88.4 | 94.5 |
| Pfly | 83.0 | 92.7 | 95.9 | 98.0 |
| DeepMSPeptide | 63.7 | 81.2 | 88.7 | 94.8 |
| PepFormer | 67.2 | 83.7 | 90.2 | 95.0 |
| DeepDetect | 78.2 | 91.4 | 95.1 | 97.7 |

**Table S5**. Partitions of the Sinitcyn *et al.* test dataset[4], showing the number of flyers, non-flyers and rescored peptides that were predicted and missed by the fine-tuned (Pfly) model.

| | Number of peptides | Percentage (%) |
|---|---|---|
| Non-flyers | 16219 | 26.9 |
| Flyers | 40846 | 67.9 |
| Predicted rescored flyers | 1461 | 2.4 |
| Missed rescored flyers | 1659 | 2,8 |

**Table S6**. Performance metrics of the fine-tuned (Pfly) model on the originally labeled and rescored Sinitcyn *et al.* test dataset[4], including accuracy, precision, recall, F1 score, Matthews Correlation Coefficient (MCC), and area under the curve (AUC).

| | Accuracy | Precision | Recall | F1 score | MCC | AUC |
|---|---|---|---|---|---|---|
| With Rescoring | 0.78 | 0.87 | 0.83 | 0.85 | 0.47 | 0.82 |
| Without Rescoring | 0.79 | 0.83 | 0.86 | 0.84 | 0.35 | 0.75 |

**Table S7**. Top flyers accurately predicted (%) by the fine-tuned (Pfly) model on the originally labeled and rescored Sinitcyn *et al.* test dataset[4].

| | Top 1(%) | Top 2 (%) | Top 3 (%) | Top 5 (%) |
|---|---|---|---|---|
| With Rescoring | 95.1 | 97.8 | 98.6 | 98.9 |
| Without Rescoring | 94.0 | 97.3 | 98.3 | 98.7 |
